# Mechanistic Insights into Impaired cGAS Activation in *Staphylococcus aureus* Biofilm Environments Reveal That STING Activation via 2′3′-cGAMP Restores Macrophage Immune Responses

**DOI:** 10.64898/2026.03.30.715225

**Authors:** Elisabeth Seebach, Carlos E. Perez Cevallos, E. Rosmin Schumacher, Katharina F. Kubatzky

## Abstract

Biofilm formation is a major cause of chronic implant-related bone infections and is associated with impaired immune responses. In a previous study, we identified the cGAS–STING pathway as a potential therapeutic target, as its activation—observed in response to planktonic *Staphylococcus aureus* (SA)—was absent in the corresponding biofilm setting. The present study aimed to identify potential mechanisms underlying the lack of cGAS activation in the biofilm environment. As biofilm-derived nucleases might degrade cGAS ligands, we assessed presence and activity of micrococcal nuclease in conditioned media from planktonic and biofilm-grown SA and evaluated the impact of extracellular DNases on cGAS pathway activation in macrophages. In addition, we examined altered cGAS expression, the requirement for continuous biofilm exposure and potential downstream inhibition resulting from degradation of the cGAS product. Biofilm formation was associated with dynamic nuclease expression, and exposure to the biofilm environment led to reduced cGAS levels in macrophages, accompanied by a lack of interferon response. Exogenous cGAS activation by G3-YSD failed to restore signaling, independent of nuclease activity or continuous biofilm exposure. In contrast, supplementation with the cGAS product and STING ligand 2′3′-cGAMP fully restored interferon responses and enhanced macrophage activation, indicating that increased degradation of the second messenger in the biofilm environment is not responsible for impaired pathway activation. Similar effects observed with *Staphylococcus epidermidis* and primary macrophages suggest a broader mechanism that is not SA- or cell line–specific. In conclusion, our data provide novel mechanistic insight into biofilm-mediated impairment of cGAS–STING signaling, revealing a previously unrecognized mechanism of immune evasion in staphylococcal biofilms. These findings extend our previous work and support the therapeutic potential of targeting STING as promising strategy to restore immune responses in chronic implant-related bone infections.

**Highlights:**

- Biofilm-derived factors impair cGAS–STING pathway activation and suppress interferon responses in macrophages.
- Impaired signaling is not primarily explained by extracellular micrococcal nuclease–mediated degradation of potential cGAS ligands.
- Biofilm exposure reduces cGAS expression levels and inhibits exogenous cGAS activation independently of continuous presence.
- Exogenous 2′3′-cGAMP fully restores interferon responses, indicating that impaired signaling is not due to degradation of the cGAS product.
- Direct activation of STING broadly enhances macrophage activation and by this could amplify overall immune responses.
- Bypassing cGAS via direct STING targeting represents a potential therapeutic strategy to overcome immune evasion in chronic implant-related bone infections.

## 1. Introduction

The formation of bacterial biofilms on implant surfaces is a major cause of chronic implant-related bone infections, including prosthetic joint infections. Biofilms protect bacteria from antibiotics and host immune defenses, thereby facilitating bacterial persistence and promoting chronic disease [1, 2]. However, biofilms do not only act as a physical barrier. Increasing evidence suggests that they actively influence their surrounding microenvironment. In particular, biofilms can shape local immune responses and promote a state of reduced immune activity [3, 4], similar to mechanisms described in the tumor microenvironment [5].

Because standard treatment strategies, such as surgical revision combined with prolonged antibiotic therapy, are often insufficient to eradicate biofilm-associated infections, additional therapeutic approaches are required [5, 6]. In this context, immunomodulatory strategies aimed at restoring or enhancing host immune responses to overcome biofilm-mediated immune suppression have gained attention as an additional therapeutic pillar [5, 7, 8]. A better understanding of the immune signaling pathways involved in biofilm-associated infections is therefore essential for the development of novel treatment strategies.

Among the innate immune pathways important for host defense, the cyclic GMP–AMP synthase (cGAS)–stimulator of interferon genes (STING) pathway plays a central role in sensing cytosolic DNA and inducing type I interferon responses [9]. Upon recognition of double-stranded DNA of endogenous or foreign origin, cGAS synthesizes the second messenger 2′3′-cGAMP, which binds to and activates STING. STING activation initiates downstream signaling cascades, including phosphorylation of IRF3 and activation of NF-κB [10, 11]. These events lead to the production of interferon-β (IFN-β), as well as proinflammatory cytokines, including interleukin-6 (IL-6). Upon binding to its receptor, IFN-β activates the JAK–STAT signaling cascade, resulting in phosphorylation of STAT1 and STAT2 and subsequent formation of the ISGF3 complex (STAT1, STAT2 and IRF9), which drives the transcription of interferon-stimulated genes such as ISG15 [12].

In a previous study, we demonstrated that conditioned medium derived from planktonic *Staphylococcus aureus* (SA) cultures (PCM) induces a robust STING-dependent response in macrophages, whereas conditioned medium obtained from SA biofilm cultures (BCM) fails to elicit a comparable activation [13]. Initially, we hypothesized that differences in the production of bacterial-derived cyclic dinucleotides (CDNs) between planktonic and biofilm growth might account for this effect. However, supplementation with bacterial 3′3′-cGAMP restored STING pathway activation in the presence of BCM, indicating that the signaling machinery downstream of STING remains functional. Further analysis revealed that this response was dependent on cGAS and that cGAS activation was absent in the biofilm setting, supporting the notion that the defect occurs upstream of STING and is not due to a lack of bacterial-derived CDNs. The absence of cGAS activation could further not be explained by a lack of extracellular bacterial DNA as this was readily detectable in SA BCM [14]. Moreover, exogenous stimulation with the cGAS ligand G3-YSD failed to induce pathway activation in the biofilm context, suggesting that the biofilm environment may both lack the appropriate cGAS-activating stimulus and directly impair cGAS signaling [13]. Based on these findings, we proposed that virulence factors present in SA PCM may induce cellular stress or damage, leading to the release of host-derived DNA into the cytosol and subsequent activation of cGAS, a process that may not occur in the biofilm environment. However, the mechanisms underlying impaired cGAS activation in response to BCM remained unresolved.

Several mechanisms may account for the absence of cGAS activation in the biofilm context: First, although extracellular DNA is present or exogenously added, its availability for cGAS sensing may be limited, for example due to degradation. Notably, SA biofilms secrete micrococcal nuclease (Nuc), which degrades extracellular DNA as part of immune evasion strategies [15, 16], potentially limiting ligand availability for cGAS. Second, biofilm-associated factors may directly affect cGAS. This could involve changes in cGAS expression, protein levels or enzymatic function, thereby limiting pathway activation independently of ligand availability. Third, inhibition may occur downstream of cGAS, for example through phosphodiesterase-mediated degradation of its second messenger 2’3’-cGAMP.

Based on these considerations, the aim of the present study was to investigate the mechanisms underlying impaired cGAS activation in the biofilm context and to identify potential strategies to overcome this impairment and to restore immune responses in chronic implant-related bone infections. To address these questions, we assessed micrococcal nuclease activity in conditioned media from planktonic and biofilm-grown SA and evaluated its impact on exogenous cGAS pathway activation by G3-YSD in macrophages. In addition, we examined cGAS expression, the requirement for continuous biofilm exposure and potential downstream inhibition by assessing subsequent STING activation after adding the cGAS product 2′3′-cGAMP.

## 2. Material and Methods

### 2.1. Bacterial culture and preparation of conditioned media

Conditioned media (CM) were generated from *Staphylococcus aureus* (ATCC 49230, UAMS-1), originally isolated from a patient with chronic osteomyelitis [17] and as described before [18]. Bacteria were cultured on Columbia agar plates supplemented with 5% sheep blood (BD Biosciences, USA). For CM preparation, colonies were inoculated into tryptic soy broth (TSB; BD Biosciences, USA) and grown for 3 h at 37 °C under shaking to obtain log-phase bacteria. A total of 6 × 10⁵ CFU/ml were transferred to DMEM high glucose (Anprotec, Germany) supplemented with 10% heat-inactivated fetal calf serum (FCS; Anprotec, Germany) and cultured at 37 °C and 5% CO₂ either under continuous shaking (200 rpm) to generate planktonic cultures or under static conditions in 24-well plates to allow biofilm formation. Biofilms were maintained for 1, 3 or 6 days with daily medium exchange. During the final 24 h of day 3 or 6 biofilm cultures, media were supplemented with 100 µg/ml gentamicin solution (Sigma-Aldrich, USA) to eliminate residual growth of planktonic bacteria. Bacterial culture supernatants were collected after 24 h of planktonic growth (PCM), after 24 hours of biofilm culture or from the last 24 h interval of day 3 or day 6 biofilms (BCM) and cleared by centrifugation (4000 rpm, 15 min, 4 °C). To exclude contamination by other bacteria, aliquots were plated on agar and incubated overnight. CM were sterile-filtered (0.2 µm), adjusted to physiological pH, aliquoted and stored at −70 °C. For early planktonic-to-biofilm transition analysis, bacteria were plated accordingly into 24-well plates and cultivated under static conditions at 37 °C and 5% CO_2_ for 2, 4, 6, 20 and 24 hours for biofilm (B-SA) and for 6 and 24 hours under continuous shaking for planktonic bacteria (P-SA), respectively. DNA digestion in PCM was done by a 60 min treatment at 37 °C with DNase I (Promega Corporation, USA). Heat inactivation of BCM was done by thermal treatment at 99 °C for 10 min. *Staphylococcus epidermidis* (SE) CM were generated accordingly and as previously described [18].

### 2.2. Nuclease activity testing and agarose gel electrophoresis

To assess nuclease activity in the CM, 20 µL of three independent CM batches were each incubated with 10 µL λ-DNA (0.3 µg/µL; Thermo Fisher Scientific, USA) at 37 °C for 10 or 90 min. Reactions were terminated by immediate freezing at −20°C. To evaluate heat stability of potential nucleases, CM samples were heat-treated at 99 °C for 10 min prior to incubation with λ-DNA under the same conditions described above. For agarose gel electrophoresis, 1 µL Midori Green Direct Stain (Nippon Genetics Europe GmbH, Germany) was added to 10 µL of each reaction sample, and 5 µL of the mixture were loaded onto a 1.2% (w/v) agarose gel prepared in TAE buffer. A 1 kb DNA ladder and a 100 bp DNA ladder (both, Nippon Genetics Europe GmbH, Germany) were included as molecular weight markers. Electrophoresis was performed at 100 V for 30 min. DNA bands were visualized and documented using a UV transillumination system (ChemoStar Imaging Systems; Intas Science Imaging Instruments GmbH, Germany).

### 2.3. Cell culture and stimulation of macrophages

#### 2.3.1 RAW 264.7 cell line

Most experiments were performed using the murine macrophage cell line RAW 264.7 (ATCC TIB-71, USA) [19]. Cells were maintained in DMEM high glucose supplemented with 10% heat-inactivated FCS and 1% penicillin/streptomycin at 37 °C and 5% CO₂. Depending on the assay, cells were seeded at densities optimized for gene expression (RT-qPCR: 300.000 cells per ml in 48-well plate with 0.5 ml in total), Western blot (WB) or flow cytometer (FACS) analysis (both: 1 x 10^6^ cells per ml in 12-well plate with 1 ml total). Cells were stimulated in CM diluted 1:1 in fresh growth medium with the cGAS agonist G3-YSD or YSD-control complexed with LyoVec^TM^ (500 ng/ml) or the cGAS product 2’3’-cGAMP (10 µg/ml, both InvivoGen, USA). RT-qPCR and WB were assessed after 20 hours of stimulation, FACS analysis of surface markers was evaluated after 24 hours of stimulation. Sequential stimulation was performed with 8 hours cultivation in BCM followed by a 20 h stimulation with the agonists. For DNase I digestion experiments, G3-YSD was either treated with 1 U DNase I per 1 µg G3-YSD for 1 hour at 37 °C followed by a 10 min at 65°C inactivation with Stop Solution according to the manufacturers protocol (Promega Corporation, USA) and then transferred to LyoVec^TM^ or corresponding amount of DNase I (2.5 U in total) were directly added in the BCM together with G3-YSD already complexed with LyoVec^TM^. The synthetic TLR2/TLR1 agonist Pam3CSK4 (P3, 10 µg/ml; InvivoGen, USA) was included as a positive control in the FACS analysis.

#### 2.3.2 Primary bone marrow-derived macrophages

Bone marrow was isolated from femora and tibiae of adult C57BL/6 mice by centrifugation at 8000 rpm for 3 min. Bone marrow–derived macrophages (BMDMs) were differentiated for 7 days in primary macrophage media (DMEM high glucose + 10% FCS + 1% Pen/Strep + 50 µM beta-mercaptoethanol) with 10–30% GM-CSF and M-CSF enriched L929 cell culture medium. Experiments were performed in primary macrophage media supplemented with 25 ng/ml M-CSF (Bio-Techne Ltd., UK). All animal procedures were conducted in accordance with national guidelines for animal care and were approved by the local authorities (T-38/24). For WB and FACS analyses, 2 × 10⁶ cells/mL were seeded into 6-well plates. Cells were stimulated with 1 ml conditioned medium in the presence or absence of the cGAS/STING agonists (G3-YSD: 500 ng/mL complexed with LyoVec; 2′3′-cGAMP: 5 µg/mL) for 20 h (WB) or 24 h (FACS). For gene expression analysis, 5 × 10⁵ cells in 0.5 mL were seeded into 24-well plates and stimulated under the same conditions for 20 h.

### 2.4. Gene Expression Analysis of Mammalian Cells and Bacteria

Total RNA from mammalian cells was isolated using the innuPREP RNA Mini Kit 2.0 (AJ Innuscreen GmbH, Germany) according to the manufacturer’s instructions. Cells were lysed in the provided lysis buffer, and lysates were applied to a DNA elimination column to remove residual genomic DNA. After addition of 70% ethanol, RNA was transferred to an RNA binding column, washed and eluted in RNase-free H₂O.

For bacterial samples, planktonic bacteria were harvested by centrifugation of liquid cultures. Biofilm-associated bacteria were collected by carefully removing supernatants, mechanically detaching the biofilm matrix from the well bottom, and centrifuging the pooled material. Bacterial pellets were stored dry at −70°C. After thawing, pellets were resuspended in TE buffer and subjected to mechanical disruption by sonication (SONOREX SUPER RK 102 H ultrasonic bath, BANDELIN electronic GmbH & Co. KG, Germany). Enzymatic lysis was performed using the innuPREP Bacteria Lysis Booster (AJ Innuscreen GmbH, Germany), and total RNA was isolated according to the manufacturer’s protocol and described above. Residual genomic DNA was removed by DNase I treatment prior to cDNA synthesis according to the manufacturer’s instructions (Promega Corporation, USA).

Complementary DNA (cDNA) was synthesized from 1 µg of total RNA using the RevertUP™ II cDNA Synthesis Kit (Biotechrabbit GmbH, Germany) according to manufacturer’s recommendations with random hexamer primers; for mammalian samples, a combination of oligo(dT) and random hexamer primers was used. A no-reverse transcriptase control (noRT) was included to exclude genomic DNA contamination. Quantitative PCR was performed using 2.5 µL of 1:2.5 diluted cDNA in a two-step amplification protocol on a StepOnePlus Real-Time PCR System (Applied Biosystems, USA) with 2× qPCRBIO SyGreen Mix Hi-ROX (PCR Biosystems Ltd., UK). Amplification specificity was confirmed by melting curve analysis and comparison with noRT and no-template (water) controls. Relative mRNA expression levels were calculated using the 2^−ΔCq method. Expression levels were normalized to *Hprt1* in mouse cells and to *gyrB* in SA samples. Mouse primers were designed using Primer3 software (https://primer3.ut.ee) based on murine gene sequences retrieved from the Ensembl genome browser (https://www.ensembl.org). Whenever possible, primers were designed to span exon–exon junctions and to detect all known transcript variants. Primer specificity was validated using NCBI Primer-BLAST (https://www.ncbi.nlm.nih.gov/tools/primer-blast). *Staphylococcus aureus*-specific primers were designed as previously described [13]. Genome sequences were retrieved from the Ensembl Bacteria genome browser (http://bacteria.ensembl.org/index.html) based on the reference genome sequence of *Staphylococcus aureus subsp. aureus USA300_FPR3757* (GCA_000013465). Primer specificity was verified in silico. All primers were synthesized by Biomers.net GmbH (Germany) and primer sequences are listed in Table 1 for mouse and Table 2 for SA.

**Table 1.**
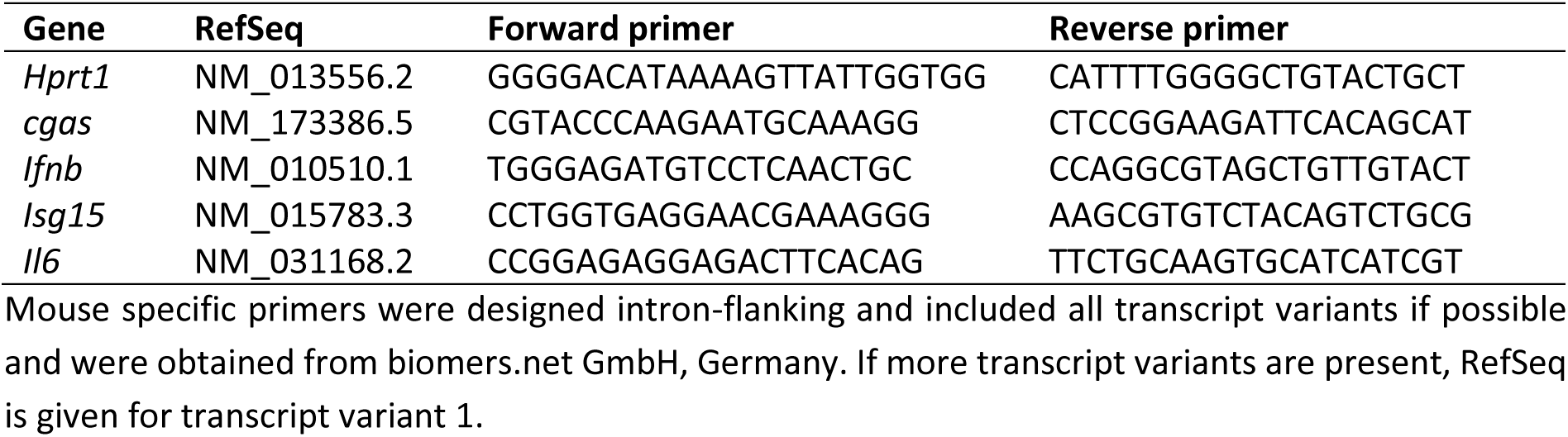
List of mouse specific oligonucleotides used for quantitative RT-PCR analysis.

**Table 2:**
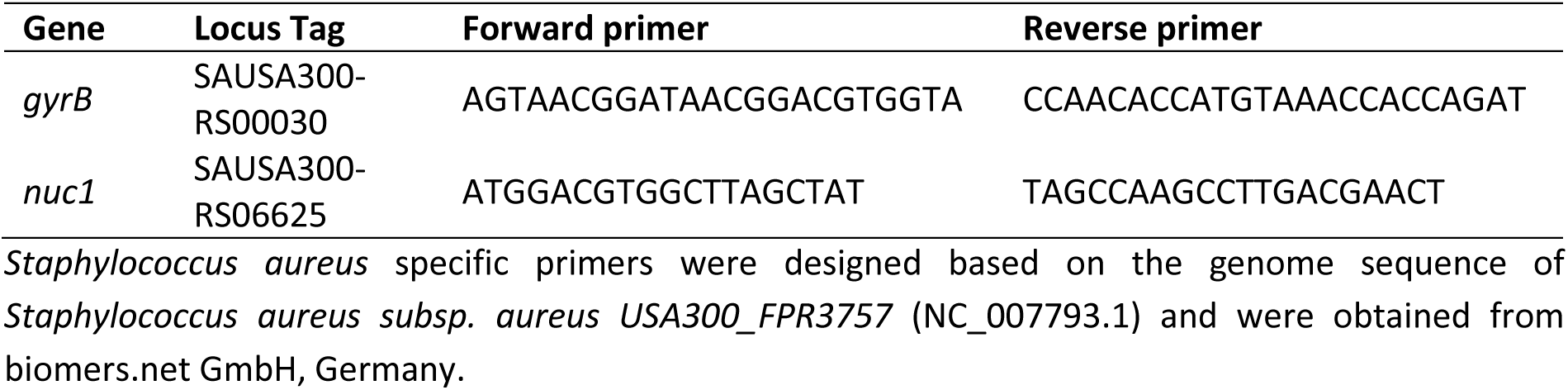
List of SA specific oligonucleotides used for quantitative RT-PCR analysis.

### 2.5. Immunoblotting

For Western blot analysis, cells were lysed in RIPA buffer (1% v/v NP-40 (Igepal CA-630), 0.25% sodium deoxycholate, 50 mM Tris-HCl pH 8.0, 150 mM NaCl, 1 mM EDTA pH 8.0, 1 mM Na₃VO₄) supplemented with protease inhibitor mix (Protease Inhibitor Mix G) and phosphatase inhibitor mix (Phosphatase Inhibitor Mix II; both Serva Electrophoresis GmbH, Germany). Lysis was performed for 1 h at 4 °C under continuous rotation. Lysates were centrifuged, and protein concentrations were determined using a BCA assay (Cyanagen Srl, Italy). Equal amounts of protein (10 µg) were mixed with Laemmli sample buffer (Serva Electrophoresis GmbH, Germany), loaded onto pre-cast 4–20% Tris-glycine gradient gels (anamed Elektrophorese GmbH, Germany), and separated by SDS-PAGE at 120 V. A broad-range, prestained protein ladder (New England Biolabs GmbH, Germany) was included on each gel. Proteins were transferred onto Amersham™ Protran™ 0.45 µm nitrocellulose membranes (GE Healthcare Life Sciences, UK). Membranes were assembled with four Whatman filter papers on each side and soaked in transfer buffer (192 mM glycine, 25 mM Tris, 2.6 mM SDS, 0.5 mM Na₃VO₄, 15% v/v methanol). Transfer was carried out for 1 h at 2 mA/cm². Successful protein transfer was verified by Ponceau S staining (0.5% Ponceau S, 3% trichloroacetic acid in ddH₂O). Membranes were blocked with BlueBlock PF (Serva Electrophoresis GmbH, Germany) with 0.5 mM Na_3_VO_4_ for 30 min at room temperature and incubated overnight at 4 °C with primary antibodies diluted in BlueBlock PF. Primary antibodies used for immunoblotting are listed in Table 3. After washing, membranes were incubated with the appropriate horseradish peroxidase (HRP)-conjugated secondary antibodies for 1 h at room temperature. Protein bands were detected using an enhanced chemiluminescence (ECL) substrate (WESTAR ETA C ULTRA 2.0, Cyanagen Srl, Italy) and visualized with a ChemoStar ECL and Fluorescence Imager (Intas Science Imaging Instruments GmbH, Germany). Band intensities were quantified using LabImage software (INTAS Science Imaging Instruments GmbH, Germany).

**Table 3.** List of antibodies used for Immunoblotting (Western Blot).

| Protein | Source | Size (kDa) | Dilution | Company |
| --- | --- | --- | --- | --- |
| Phospho-STAT1 (Tyr701) | rabbit | 84, 91 | 1:1000 | Cell Signaling Technology, USA |
| cGAS | rabbit | 62 | 1:1000 | Cell Signaling Technology, USA |
| β-Actin | rabbit | 42 | 1:1000 | Proteintech, USA |
| GAPDH | mouse | 36 | 1:1000 | Proteintech, USA |
| Anti-mouse IgG, HRP-linked | goat |  | 1:1000 | Cell Signaling Technology, USA |
| Anti-rabbit IgG, HRP-linked | goat |  | 1:1000 | Cell Signaling Technology, USA |

Antibodies were all recommended for use in mouse and applied according to the manufacturer's advice. Proteins were detected by chemiluminescent luminol reaction after incubation with respective HRP-linked secondary antibody and imaged in a ChemoStar ECL Imager.

### 2.6. Flow Cytometry

Supernatants were collected and stored at −70 °C. Cells were washed twice with PBS, scraped in PBS and centrifuged at 5000 rpm for 2 min. Cells were resuspended in PBS containing 2% BSA. Unstained controls were incubated in PBS/2% BSA alone. For antibody staining, 0.2 mg/mL FITC-conjugated anti-TLR2 (Novus Biologicals, UK), PE-conjugated anti-MHC II (Invitrogen, USA), PE-conjugated anti-CD80, or PE-conjugated anti-CD86 (both BioLegend, USA) were added and cells were incubated for 1 h at 4 °C in the dark. Following incubation, cells were washed twice with PBS, resuspended in PBS and analyzed using a BD FACSCanto™ flow cytometer and BD FACSDiva™ Software (BD Biosciences, USA).

### 2.7. Cytometric Bead Array

Cell culture supernatants obtained from experiments performed for FACS analysis were used for cytokine quantification by cytometric bead array (CBA; LEGENDplex™, BioLegend, USA). The assay was conducted according to the manufacturer’s instructions. Briefly, a Mouse Inflammation Panel (Mix and Match Subpanel) comprising beads specific for IFN-β and IL-6 was employed. After centrifugation, supernatants were diluted 1:5 in Assay Buffer and transferred to a V-bottom 96-well plate. A standard curve was included in the experiment. Following the addition of the bead mixture, samples were incubated overnight at 4 °C in the dark under continuous shaking. After washing steps, detection antibodies were added and incubated for 1 h at room temperature. Subsequently, streptavidin-phycoerythrin (SA-PE) was added, and samples were incubated for an additional 30 min at room temperature with shaking. Samples were acquired using a MACSQuant® 10 flow cytometer (Miltenyi Biotec, Germany). Cytokine concentrations were calculated using the LEGENDplex™ Data Analysis Software (https://legendplex.qognit.com) according to the manufacturer’s guidelines.

### 2.8. Statistical Analysis

Unless otherwise stated in the figure legends, all experiments were performed with three independent biological replicates. Data are presented as mean ± standard error of the mean (SEM), with individual data points shown as dots. Statistical analyses were conducted using an ordinary one-way analysis of variance (ANOVA) followed by post hoc multiple comparison testing with Bonferroni correction. A p-value < 0.05 was considered statistically significant. All statistical analyses were performed using GraphPad Prism 10 for Windows (Version 10.6.1; GraphPad Software Inc., USA).

## 3. Results

### SA biofilm formation is accompanied by pronounced thermonuclease activity in the biofilm microenvironment, particularly during the planktonic-to-biofilm transition

In contrast to our previous studies, gentamicin treatment was incorporated into the biofilm protocol to exclude potential contributions of planktonic bacteria to SA BCM. Therefore, gentamicin was added during the final medium exchange prior to collection of biofilm supernatants. To verify biofilm formation and the effectiveness of gentamicin treatment, crystal violet staining and outgrowth cultures of biofilms and corresponding supernatants were performed. These analyses confirmed the formation of a robust biofilm after six days of culture. Notably, bacterial growth remained detectable from biofilm-derived streak-outs despite gentamicin treatment, indicating persistence of biofilm-associated SA. In contrast, no bacterial growth was observed in supernatants collected from gentamicin-containing cultures, confirming effective elimination of planktonic bacteria (Fig. 1A). Progressive acidification of the microenvironment was observed over the course of biofilm formation, as indicated by a color change of the phenol red indicator in the culture medium from purple to yellow (Fig. 1B). This was likely due to increased lactate production by biofilm-associated bacteria [18]. In three-day-old biofilms, gentamicin treatment was associated with a reduced metabolic turnover rate. This difference was no longer apparent after six days. At this later time point, both gentamicin treated and untreated biofilms exhibited a highly acidified environment. Since acidification and lactate accumulation in the microenvironment are hallmarks of mature biofilms, day 6 BCM generated under gentamicin treatment was used for subsequent experiments to exclude potential contributions from planktonic bacteria. All CM exhibited substantial nuclease activity, as demonstrated by efficient digestion of λ DNA (Fig. 1C). Although gentamicin treatment was associated with reduced nuclease activity, a considerable capacity to degrade λ DNA remained. Remarkably, the highest nuclease activity was detected in day 1 BCM, where complete degradation of λ DNA was observed within 90 minutes. Heat inactivation of CM did not abolish nuclease activity, consistent with the known heat resistance of the SA thermonuclease Nuc [20]. Comparable results were obtained for the MRSA strain USA300 (Fig. 1D). In contrast, SE does not possess a thermostable nuclease [21]. Accordingly, SE CM displayed lower nuclease activity compared to SA CM, and this activity was completely abrogated after heat inactivation (Fig. 1E). To further investigate nuclease regulation, *nuc1* gene expression was analyzed during biofilm formation. The highest mRNA levels were detected in planktonic bacteria. Expression decreased after 24 hours of biofilm formation and subsequently increased again at later time points (Fig. 1F left). *Nuc1* expression was not affected by gentamicin treatment (Fig. 1F right). Since these expression data did not fully correspond to the nuclease activity assay—where maximal activity was observed in day 1 BCM despite relatively low *nuc1* expression in day 1 biofilm samples—we examined earlier stages of biofilm development. Indeed, *nuc1* mRNA levels exhibited a pronounced peak after six hours of static SA cultivation (Fig. 1G, left), corresponding to the adhesion and transition phase from planktonic to biofilm growth [22]. This peak was not observed under continuous shaking conditions, where bacteria were maintained in a planktonic state (Fig. 1G, right). After 24 hours, *nuc1* expression dropped in the biofilm culture, whereas it increased slightly in the planktonic culture.

**Figure 1.**
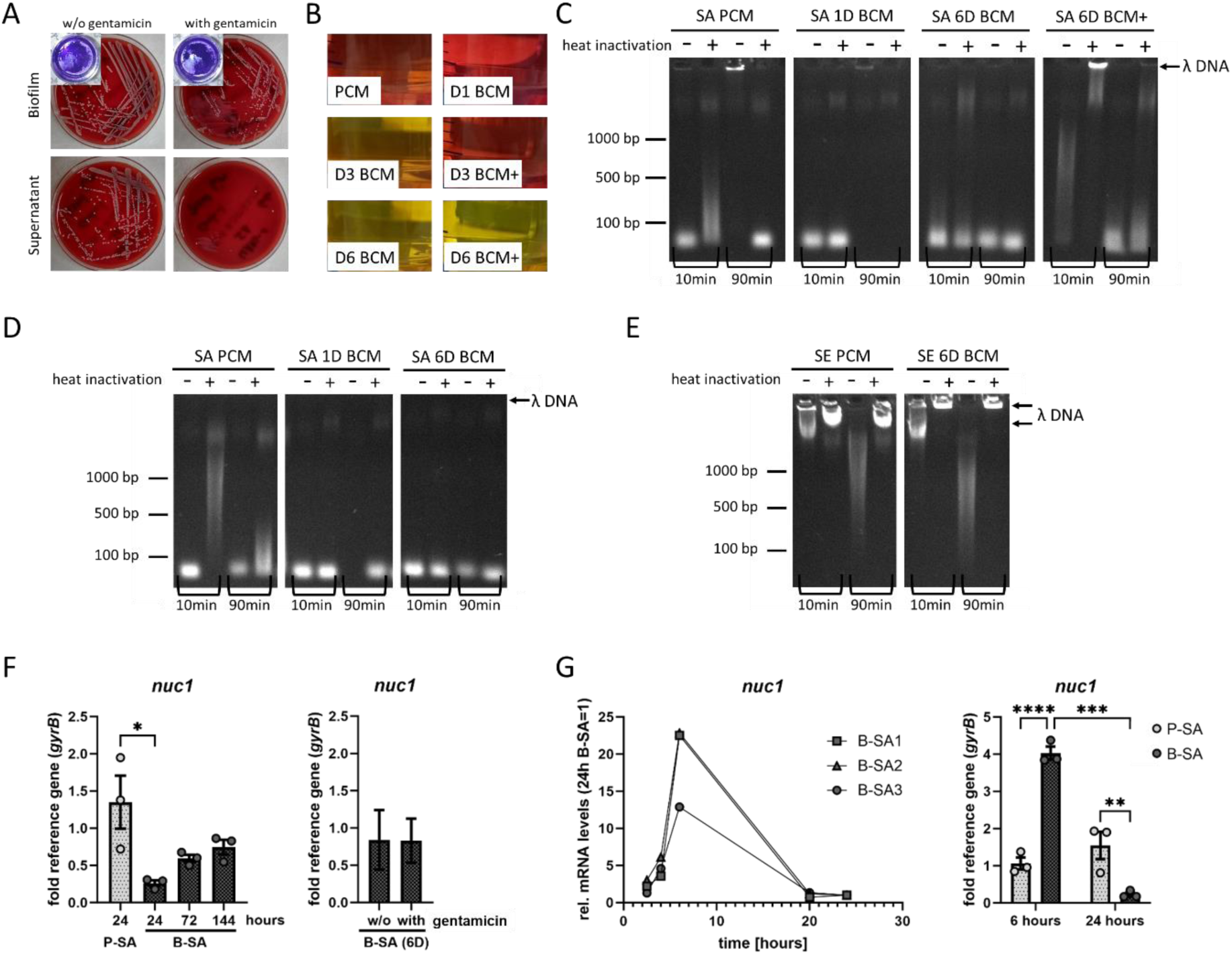
Characterization of the CM. *Staphylococcus aureus* UAMS-1 was cultured in DMEM high glucose + 10% heat-inactivated FCS either under continuous shaking for planktonic growth or under static conditions in cell-culture treated well-plates for biofilm formation with daily medium exchange. CM were harvested directly after the first 24 hours for PCM and 1D BCM or 24 hours after medium exchange at day 3 or day 6 of the biofilm culture. D3 and D6 BCM were cultured either with or without the addition of 100 µg/ml gentamicin to suppress planktonic growth. A) Biofilm formation after 6 days of culture with or without gentamicin after Crystal Violet staining is shown in the small pictures. Bacteria were streaked out on blood agar plates either from 6D biofilms (upper pictures) or the 6D biofilm supernatants (lower pictures) to assess bacterial growth and to confirm purity, ensuring that no contaminants other than the inoculated SA strain were present. B) Representative images showing the appearance of CM obtained at different culture times (1D, 3D and 6D) and under various conditions (+ indicates gentamicin treatment) after sterile filtration and prior to pH adjustment. C-E) Representative agarose gels showing nuclease activity as evidenced by λ DNA digestion after incubation in different CM of SA strain UAMS-1 (C), SA strain USA300 (D) or *Staphylococcus epidermidis* (SE) strain RP62A (E) for 10 or 90 min. CM were used directly or after heat inactivation at 99 °C for 10 minutes. F+G) Expression levels of the micrococcal nuclease gene *nuc1* during biofilm formation (F) and the planktonic-to-biofilm transition phase (G). Planktonic (P-SA) and biofilm-grown (B-SA) SA were harvested at the indicated time points. *Nuc1* mRNA levels were quantified by RT-qPCR and normalized to the reference gene *gyrB*. Graphs show mean ± SEM, with independent experiments indicated as dots. Statistical comparisons were performed using one-way ANOVA followed by Bonferroni’s post hoc test. P < 0.05 was considered statistically significant.

### Differential interferon responses to SA PCM and BCM are associated with reduced cGAS expression and extracellular DNA

As previously reported, SA PCM induce a pronounced cGAS-STING–IRF3-dependent interferon response, whereas this response was absent in the corresponding biofilm setting [13]. Here, we aimed to (i) validate this finding using SA BCM generated under gentamicin treatment to exclude contributions from planktonic bacteria, (ii) rule out a potential bias of gentamicin itself on the macrophage response, and (iii) investigate cGAS expression as a potential mechanism underlying impaired pathway activation in the biofilm context. *Cgas* mRNA levels were slightly reduced in BCM compared to PCM. Consistent with our previous findings, PCM—but not BCM—induced *Ifnb* and *Isg15* gene expression, whereas both PCM and BCM led to increased *Il6* expression (Fig. 2A). This pattern was also reflected at the protein level. Increased cGAS protein levels and STAT1 phosphorylation were observed following IFN-β and PCM stimulation, whereas cGAS protein abundance was markedly reduced in cells exposed to BCM (Fig. 2B,C). Given the previously observed differences in extracellular DNA levels between planktonic and biofilm-conditioned media [14], we next asked whether cGAS activation in PCM depends on extracellular DNA. Therefore, PCM was treated with DNase I prior to macrophage stimulation. DNase I digestion resulted in a moderate reduction in *cgas* mRNA levels, the interferon response, and *Il6* expression (Fig. 2D). However, these levels remained substantially higher than those observed in BCM-treated cells. Interestingly, as previously reported, compared to SA, SE PCM showed high levels of extracellular DNA and also induced an interferon response; however, in an IRF3- and thus cGAS-STING-independent manner [14, 18]. Accordingly, increased p-STAT1 protein levels were detected in PCM-treated cells (Fig. 2E), whereas cGAS expression remained low on protein and mRNA level (Fig. 2E,F). DNase I digestion of SE PCM resulted in a slightly more pronounced reduction of the interferon response (Fig. 2F), indicating that extracellular DNA contributes to pathway activation in both conditions although likely through distinct recognition mechanisms, as previously suggested.

**Figure 2.**
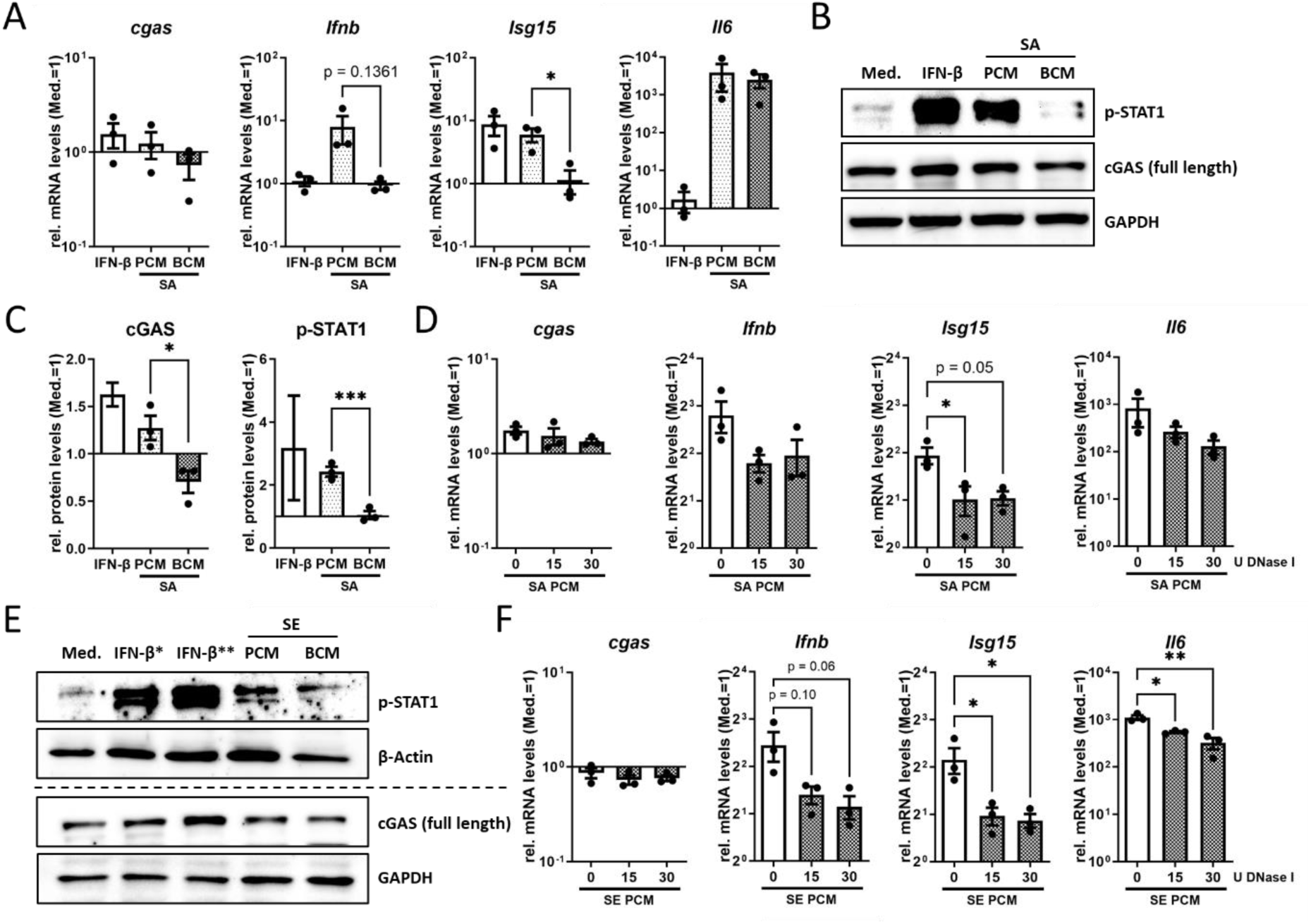
cGAS-STING pathway activation and type I IFN response induced by CM. RAW 264.7 cells were treated with CM diluted 1:1 in fresh culture medium for 20 h. Recombinant mouse IFN-β (*2 or **20 ng/ml; BioLegend, USA) served as a positive control. A) Gene expression levels of *cgas*, *Ifnb*, *Isg15* and *Il6* as indicators of type I IFN response and general immune activation (*Il6*) following CM treatment. mRNA levels were quantified by RT-qPCR and normalized to the reference gene *Hprt1* and expressed relative to the medium-treated control (Med.). B, C, E) Protein analysis by Western blot. (B) Representative WB image; bands were cropped for presentation, and the full blot is shown in Supplementary Fig. 1. (C) Quantification of three independent blots. Protein levels were normalized to the loading control GAPDH and expressed relative to the medium-treated control (Med.). (E) Representative WB image after treatment of cells with CM generated from SE RP62A. D+F) Contribution of extracellular DNA in PCM to the induction of a type I IFN response. PCM of SA (D) or SE (F) were treated with DNAse I at 37 °C for 1 h. Cells were then treated with these media for 20 h. mRNA levels of *cgas*, *Ifnb*, *Isg15* and *Il6* were assessed by RT-qPCR analysis, normalized to the reference gene *Hprt1* and expressed relative to the medium-treated control (Med.). Graphs show mean ± SEM, with independent experiments indicated as dots. Statistical comparisons were performed using one-way ANOVA followed by Bonferroni’s post hoc test. P < 0.05 was considered statistically significant.

### SA BCM impairs exogenous cGAS activation independently of nuclease activity and persistent BCM exposure

As our previous data indicated that exogenous cGAS activation by G3-YSD is impaired in the presence of SA BCM, we next aimed to validate this finding in our modified experimental setting. To ensure that agonist concentration was not limiting, we stimulated RAW 264.7 cells with a fivefold higher concentration of G3-YSD (500 ng/ml vs. 100 ng/ml previously [13]). Consistent with our previous findings, gene expression analysis revealed that no interferon response was detectable in the presence of BCM (Fig. 3A). To exclude a potential effect of residual gentamicin in BCM, we assessed interferon induction upon direct addition of gentamicin during G3-YSD stimulation in normal cell culture medium. Although fresh gentamicin slightly reduced gene expression levels, residual antibiotic in BCM could be ruled out as the cause of impaired cGAS activation (Suppl. Fig. 2A). Finally, to determine whether this effect is dependent on altered cGAS function in the transformed cell line used, we performed the same experiments in primary macrophages. The inhibitory effect of BCM was still detectable, albeit less pronounced, in BMDMs. In these cells, G3-YSD stimulation in the presence of BCM induced a delayed interferon response, characterized by an early increase in *Ifnb* expression followed by reduced induction of downstream genes such as *cgas* and *Isg15* (Fig. 3B), indicating partial cell type–specific differences.

**Figure 3.**
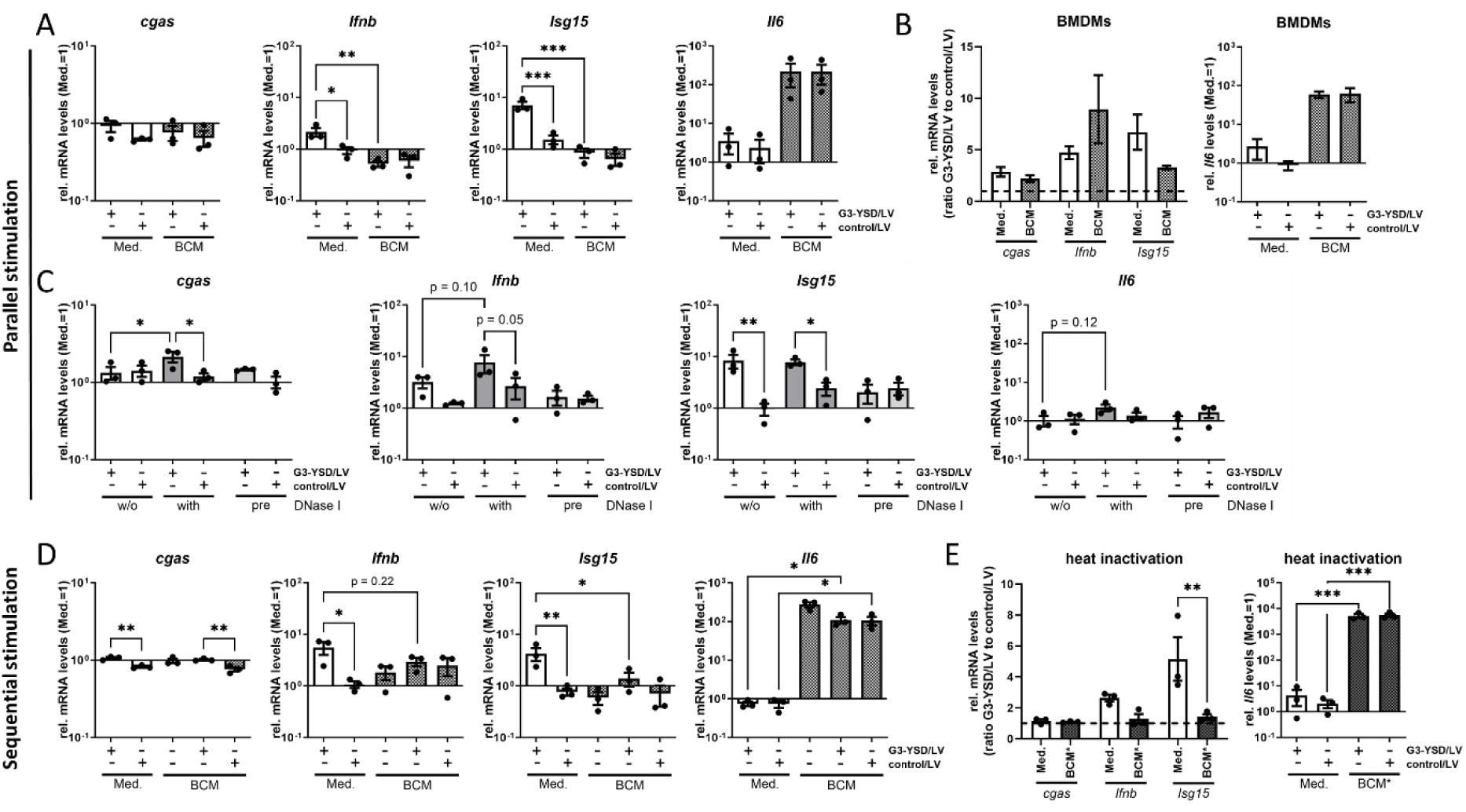
Exogenous activation of the cGAS-STING pathway in BCM and the role of nucleases. A–B) Macrophages (A: RAW 264.7 cells; B: BMDMs, n=2 mice) were treated with 500 ng/ml G3-YSD complexed with LyoVec™ in the presence or absence of SA BCM (1:1 diluted in fresh culture medium) for 20 h. C) RAW 264.7 cells were stimulated with G3-YSD complexed with LyoVec™ for 20 h. G3-YSD was either pre-digested with DNase I for 1 h at 37 °C prior to complexing with LyoVec™, or DNase I was added simultaneously with the G3-YSD/LyoVec™ complex. D) RAW 264.7 cells were pre-treated with SA BCM for 8 h, after which BCM was removed, and cells were stimulated with 500 ng/ml G3-YSD/LyoVec™ for an additional 20 h in cell culture medium. Induction of a type I IFN response and general immune activation was assessed by RT-qPCR analysis of *cgas*, *Ifnb*, *Isg15* and *Il6*. mRNA levels were normalized to the reference gene *Hprt1* and expressed relative to the medium-treated control (Med.). E) SA BCM was heat-inactivated at 99 °C for 10 min (BCM*) and subsequently added to RAW 264.7 cells together with G3-YSD/LyoVec™ or its control. Induction of the type I IFN response was assessed by RT-qPCR. Expression of *cgas*, *Ifnb* and *Isg15* is shown as the ratio of cells treated with G3-YSD to cells treated with the control in the respective medium (Med. or BCM, left graph). *Il6* gene expression was assessed as a marker of general immune activation and is shown relative to the medium-treated control (Med., right graph). Graphs show mean ± SEM, with independent experiments indicated as dots. Statistical comparisons were performed using one-way ANOVA followed by Bonferroni’s post hoc test. P < 0.05 was considered statistically significant.

To assess whether nuclease activity contributes to the lack of G3-YSD-mediated signaling in BCM through degradation of the DNA, DNase I digestion experiments were performed. DNase I treatment of G3-YSD prior to complexing with the transfection reagent LyoVec^TM^ efficiently reduced the interferon response, confirming that DNase I digestion was effective (Fig. 3C). In contrast, the addition of DNase I together with G3-YSD/LyoVec^TM^ did not diminish the interferon response but rather increased *cgas* and *Ifnb* expression. These findings indicate that nuclease activity is unlikely to account for the absence of cGAS activation by G3-YSD in SA BCM. We therefore investigated whether the continuous presence of BCM is required for the impairment of exogenous cGAS-STING pathway activation. RAW 264.7 cells were pretreated with BCM for 8 hours, after which the medium was removed and the cells were subsequently stimulated with G3-YSD. Under these conditions, the impairment of G3-YSD mediated cGAS-STING pathway activation persisted (Fig. 3D), indicating that the effect is long-lasting and independent of the simultaneous presence of BCM. To further evaluate whether proteins other than thermonucleases might contribute to this effect, RAW 264.7 cells were stimulated with heat-inactivated BCM. Even after heat inactivation, G3-YSD stimulation failed to induce an interferon response in the presence of BCM (Fig. 3E), suggesting that heat-stable biofilm-derived factors may contribute to this effect. In contrast, induction of *Il6* gene expression remained unaffected. Interestingly, heat-inactivated SA BCM itself seemed to result in a low and cGAS-independent induction of *Ifnb* and *Isg15* gene expression (Suppl. Fig. 2B).

### Exogenous 2′3′-cGAMP reveals that impaired signaling is not due to degradation of the cGAS product and enhances macrophage immune response in SA BCM

Degradation of the cGAS product by phosphodiesterases may represent a potential mechanism underlying impaired downstream signaling. Therefore, we next evaluated whether supplementation with 2′3′-cGAMP is sufficient to restore pathway activation in the presence of BCM. While the suppressive effect of SA BCM on *cgas* expression persisted following stimulation with 2′3′-cGAMP, *Ifnb* and *Isg15* expression were strongly induced when 2’3’-cGAMP was added to BCM, exceeding the levels observed with 2’3’-cGAMP stimulation alone (Fig. 4A, for *Isg15*: cGAMP alone versus cGAMP in BCM, p=0.07). Remarkably, this effect was not limited to interferon-associated genes. *Il6* expression was also significantly increased at both the mRNA and protein level upon simultaneous stimulation with BCM and 2’3’-cGAMP (Fig. 4A,B). Similar results were obtained in primary BMDMs, confirming that this effect was not restricted to the RAW 264.7 macrophage cell line (Fig. 4C). Encouraged by the observation that both interferon-related genes and the broader inflammatory marker gene *Il6* were enhanced, we further investigated whether overall macrophage activation was increased under these conditions. Indeed, stimulation with 2’3’-cGAMP in the presence of SA BCM not only restored but further enhanced macrophage activation, as indicated by highly increased surface expression of TLR2, MHC II, CD80, and CD86 (Fig. 4D). A comparable enhancement of macrophage activation by 2’3’-cGAMP was also observed in the presence of SE BCM (Fig. 4E,F). Enhanced macrophage immune activation by 2’3’-cGAMP in SA BCM was also reproduced in BMDMs as shown by elevated CD86 levels (Suppl. Fig. 2C) and increased acidification of the medium (Suppl. Fig. 2D) indicating a more pronounced shift towards glycolysis and a pro-inflammatory macrophage polarization. Importantly, this potentiating effect of BCM on 2’3’-cGAMP mediated interferon responses required simultaneous stimulation, as it was lost when cells were first exposed to BCM followed by subsequent 2’3’-cGAMP treatment. In contrast, the increase in *Il6* gene expression remained unaffected under sequential stimulation conditions (Fig. 4F–H). Notably, even after prior exposure to BCM, induction of an interferon response by 2’3’-cGAMP remained possible.

**Figure 4.**
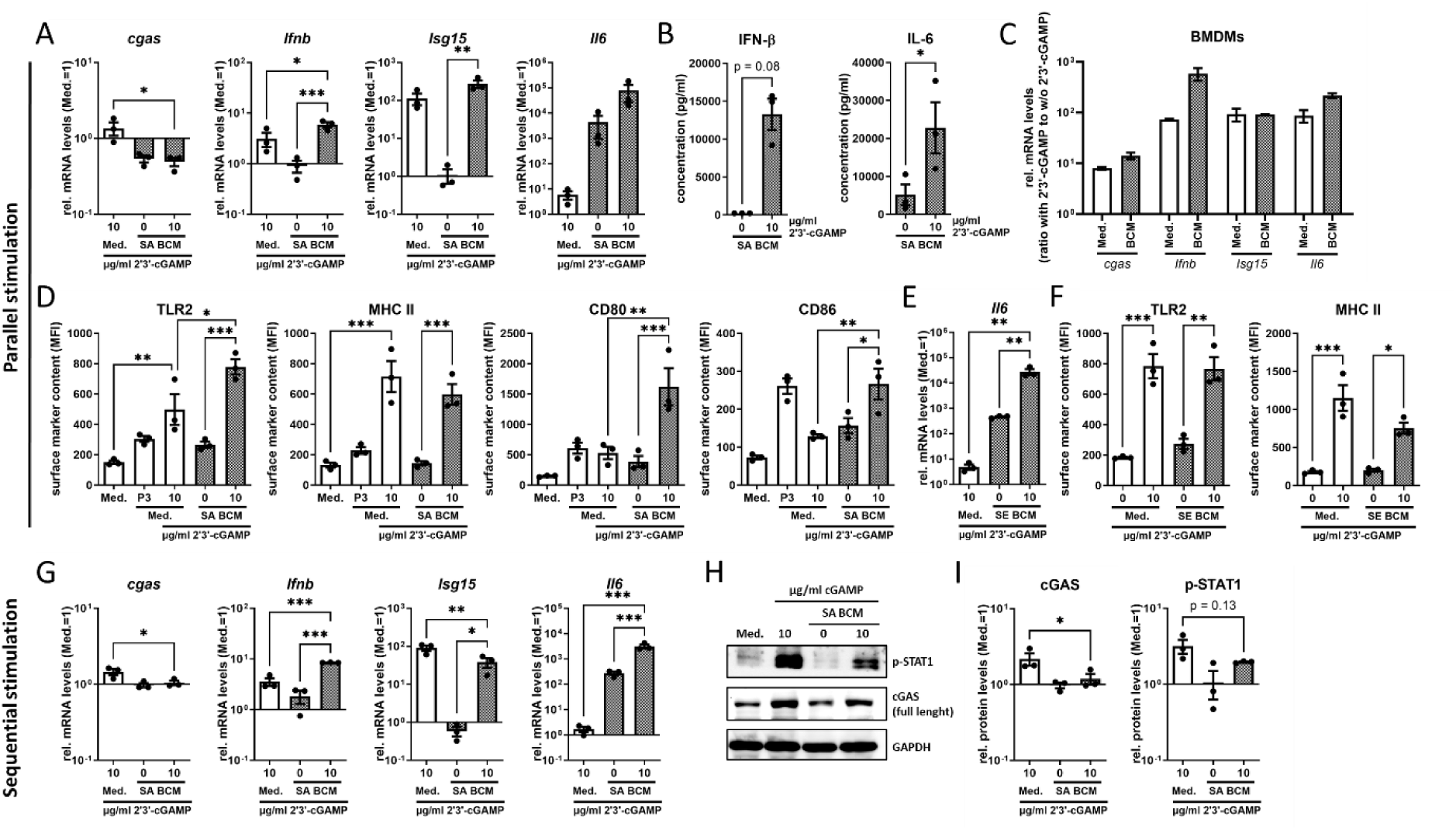
Effect of exogenous addition of the cGAS product 2’3’-cGAMP on cGAS-STING pathway activation in BCM. A–F) Macrophages (A+B/D-F: RAW 264.7 cells; C: BMDMs, n=2 mice) were treated with 5 µg/ml (BMDMs) or 10 µg/ml (RAW 264.7 cells) 2’3’-cGAMP in the presence or absence of BCM (1:1 diluted in fresh culture medium) for 20 h (RT-qPCR) or 24 h (FACS). BCM was derived from SA (A-D) or SE cultures (E+F). G-I) RAW 264.7 cells were pre-treated with SA BCM for 8 h. After removal of BCM, cells were stimulated with 10 µg/ml 2’3’-cGAMP for an additional 20 h. A, C, E, G) Induction of a type I IFN response and general immune activation was assessed by RT-qPCR analysis of *cgas*, *Ifnb*, *Isg15* and *Il6*. mRNA levels were normalized to the reference gene *Hprt1* and expressed relative to the medium-treated control (Med.). B, D, F, H, I) Protein analysis was performed by LegendPlex™ assay (B), flow cytometric analysis of surface markers (FACS; D, F), and Western blot (H, I). (H) Representative WB image; (I) quantification of three independent blots. Protein levels were normalized to the loading control GAPDH and expressed relative to the medium-treated control (Med.). Graphs show mean ± SEM, with independent experiments indicated as dots. Statistical comparisons were performed using one-way ANOVA followed by Bonferroni’s post hoc test. P < 0.05 was considered statistically significant.

## 4. Discussion

Biofilms on implant surfaces make implant-related bone infections such as periprosthetic joint infection particularly difficult to treat. Beyond the protective shielding provided by the biofilm’s extracellular polymeric substances, biofilms actively modulate their local microenvironment toward a more anti-inflammatory and immunosuppressed state. Consequently, reactivation and strengthening of an effective immune response represent promising therapeutic strategies [5, 7]. Our previous findings indicate that activation of the cGAS-STING signaling pathway is lacking in the biofilm context, identifying it as a potential therapeutic target [13]. In the present study, we provide insight into the mechanisms underlying impaired cGAS activation in the biofilm context and identify potential strategies to overcome this impairment and restore immune responses in chronic implant-related bone infections.

Our results confirm our previous finding that, in contrast to a planktonic SA environment, the corresponding biofilm condition failed to induce a type I IFN response. Building on this, we next investigated the underlying mechanisms of impaired pathway activation. Although cGAS expression was reduced following stimulation with biofilm CM, it remained detectable at both mRNA and protein levels, suggesting that diminished cGAS abundance may contribute to, but does not fully account for, the absence of pathway activation. Both planktonic and biofilm CM exhibited pronounced nuclease activity. Notably, *nuc1* expression was initially reduced after one day of biofilm formation, increased again over the course of maturation up to day 6 and peaked during the early transition phase from planktonic growth to initial biofilm adhesion. This was accompanied by elevated nuclease activity in day 1 BCM compared to PCM. However, supplementation of PCM with exogenous DNase did not eliminate the type I IFN response. Likewise, DNase treatment in standard culture medium did not prevent G3-YSD–induced cGAS activation. Collectively, these findings argue against nuclease-mediated extracellular DNA degradation as the primary mechanism underlying the absent cGAS-STING signaling observed in the biofilm context. Consistent with this interpretation, BCM prevented cGAS-STING pathway activation not only upon simultaneous G3-YSD stimulation but also after removal of BCM prior to sequential stimulation. The persistence of this inhibitory effect indicates that the mechanism does not depend on the continued presence of soluble extracellular factors such as nucleases but rather point toward a biofilm-mediated impairment of cellular cGAS-STING signaling. The phenomenon was still detectable but less pronounced in primary BMDMs, suggesting that the virus-induced cancer-derived RAW 264.7 cells may be more susceptible to biofilm-mediated inhibition of cGAS activation, possibly due to alterations in cGAS-STING pathway regulation associated with its tumor background. Based on our previous findings showing that addition of the bacterial STING ligand 3′3′-cGAMP restores pathway activation in the biofilm context, we concluded that the signaling cascade downstream of STING activation remains functional [13]. Notably, 3′3′-cGAMP is resistant to degradation by the host phosphodiesterase ENPP1, in contrast to the endogenous cGAS product 2′3′-cGAMP [23]. This raised the possibility that degradation of 2′3′-cGAMP by ENPP1 may represent an additional mechanism behind the limited cGAS-STING pathway activation in the biofilm environment, acting independently of cGAS function. Therefore, we next assessed whether exogenous supplementation with 2′3′-cGAMP can restore cGAS-STING signaling in the presence of BCM. Addition of 2′3′-cGAMP not only restored type I IFN induction but even resulted in an enhanced immune response. In addition to increased IFN-β levels, 2′3′-cGAMP stimulation increased IL-6 production as well as the expression of TLR2, MHC II, CD80 and CD86. This effect was consistently observed in both RAW 264.7 cells and BMDMs and was not restricted to SA BCM but also occurred in SE BCM, suggesting a broader relevance across staphylococcal biofilms. These findings are in line with and extend our previous observations, further supporting that impaired immune activation in the biofilm environment occurs upstream of STING activation, likely involving restricted ligand availability and impaired cGAS activity. Furthermore, our data demonstrate that direct STING activation with its endogenous second messenger 2′3′-cGAMP effectively bypasses this blockade and restores immune responses, thereby representing a promising therapeutic strategy.

In line with others, we detected expression of the secreted micrococcal thermonuclease Nuc in the SA strain UAMS-1 [24]. In the context of infections, micrococcal nucleases are primarily known to facilitate immune evasion by degrading neutrophil extracellular traps (NETs), thereby promoting bacterial survival and persistence [25]. In addition, nucleases have been implicated in biofilm biology, contributing to mature biofilm architecture and late-stage dispersal while inhibiting early biofilm formation [26]. Consistent with this notion, low nuclease expression has been associated with increased biofilm accumulation [27, 28]. However, nuclease activity remains important for biofilm development and bacterial persistence in implant infection models, largely due to its role as virulence factor in immune evasion [28, 29]. In line with this, we observed low nuc1 mRNA levels during early biofilm development (24 h), followed by increased expression during later stages of biofilm growth (6 days). Interestingly, we also observed a transient peak in *nuc1* expression approximately 6 h after initiation of static SA culture. A similar early induction of micrococcal nuclease expression has been described previously, with increased expression occurring around 4–6 h after the onset of biofilm formation under the control of the SaeRS two-component system. This early nuclease activity was found to promote a nuclease-dependent biofilm exodus that is critical for subsequent tower formation essential for normal biofilm development [30]. Whether this transient increase in micrococcal nuclease expression at the onset of biofilm formation indeed contributes to reduced pattern-recognition receptor (PRR)-mediated detection of bacterial DNA within the biofilm environment, additional to its established role in NET escape [15], remains to be determined. However, based on our current data, nuclease activity alone is unlikely to represent the primary mechanism underlying the impaired activation of the cGAS-STING pathway observed in the SA biofilm context. Accordingly, we examined whether other factors present in the SA biofilm environment contribute to this effect. Notably, the observed effect persisted even after removal of the SA BCM, suggesting that it induces cellular changes rather than requiring continuous presence of the bacterial factor. Moreover, heat inactivation of the BCM did not abolish the effect, indicating that the responsible factor is heat-stable. cGAS activation can be regulated at both the transcriptional and post-transcriptional level [31]. A role for microRNAs (miRNAs) in the regulation of cGAS expression has already been described in non-bacterial disease models [32]. In line with this, we observed a downregulation of cGAS mRNA and protein levels in the SA BCM setting. This reduction could potentially be mediated by infection-modulated miRNAs, as previously discussed in other bacterial host–pathogen interactions [33]. Furthermore, cGAS activity is tightly regulated by post-translational modifications (PTMs), including phosphorylation, ubiquitination, acetylation, glutamylation and sumoylation, many of which have been reported to inhibit cGAS activity [31, 34]. In addition, lactylation of cGAS was recently described as a modification that suppresses its activity [35, 36] and promotes its degradation [37]. Biofilm environments are typically characterized by glucose depletion and the accumulation of lactate, which is also reflected in our SA BCM [18] and by others [38]. Under such metabolic conditions, elevated extracellular lactate levels could promote increased protein lactylation. Beyond changes in levels of carbohydrates and organic acid metabolites, the amino acid and lipid composition of the biofilm environment might also be altered and may further modulate cGAS activity [39].

In contrast to the impaired cGAS activation observed in SA biofilm environment, planktonic SA environment induced a cGAS-dependent immune response. These differences highlight the profound impact of bacterial growth state on host innate immune sensing. Investigating the mechanisms underlying cGAS activation during planktonic infection may therefore provide important clues to better understand why this pathway remains inactive in the biofilm context. One potential explanation may lie in the distinct metabolic programs induced in host cells. Whereas planktonic SA infections are typically associated with glycolysis and ACOD1/itaconate-mediated breakpoints in the tricarboxylic acid (TCA) cycle, the metabolic environment created by SA biofilms has been reported to drive host cell metabolism towards increased TCA cycle activity and oxidative phosphorylation (OxPhos) [40, 41]. Dysregulated activity of the TCA cycle enzymes succinate dehydrogenase (SDH) and fumarate hydratase (FH), as well as the accumulation of fumarate and succinate, have been associated with mitochondrial stress and the release of mitochondrial RNA and DNA subsequently inducing cGAS-dependent and -independent type I interferon responses [42, 43]. In contrast, a functional TCA cycle, as suggested in the SA biofilm context, may limit inflammation-induced mitochondrial damage and thereby reduce cGAS activation triggered by the release of mitochondrial nucleic acids. Thus, increased TCA cycle activity may preserve mitochondrial integrity and represent an additional regulatory mechanism protecting against mitochondrial damage–mediated activation of the cGAS-STING pathway in the biofilm environment.

Interestingly, we found extracellular DNA to contribute to the induction of a type I interferon response in both SA and SE planktonic environments. In our previous work, this response was shown to be mediated via the cGAS–STING–IRF3 axis in SA, whereas in SE it occurred through an IRF7-dependent and therefore cGAS-independent mechanism [13, 18]. These findings suggest that bacterial DNA in the environment can engage distinct immune sensing pathways depending on the bacterial species. In the case of SE, bacterial DNA appears to be predominantly detected after phagosomal uptake via TLR9, while in SA cytosolic DNA sensing through cGAS is the main mechanism behind the type I interferon induction. How extracellular bacterial DNA present in SA PCM gains access to the host cytosol remains to be elucidated, although toxin-mediated disruption of phagosomal membranes represents a plausible mechanism. Notably, we also detected high levels of extracellular bacterial DNA in SA BCM [14]. However, in contrast to the planktonic context, this DNA did not appear to induce cGAS activation, which is consistent with our observations following exogenous DNA stimulation using G3-YSD. It should also be considered that DNase I remained present in PCM after the 1 h pre-digestion step. Thus, we cannot fully exclude the possibility that subsequent release of host cell DNA, for example as a consequence of virulence factor–mediated cellular damage induced by SA PCM, may contribute to cGAS activation and could have been degraded during DNase treatment independently of bacterial DNA. Taken together, our data support the hypothesis that the reduced virulence of SA BCM compared with PCM results in diminished host cell damage and inflammation. This, in turn, may limit the damage-associated release of host-derived nucleic acids and thereby protect against activation of innate immune sensing pathways, including the cGAS–STING axis.

Importantly, the suppressed innate immune activation observed in the biofilm context could be overcome by direct stimulation of STING using exogenous 2′3′-cGAMP, the endogenous second messenger produced by cGAS [9, 44]. The ability of cGAMP to enhance innate immune signaling is consistent with previous studies demonstrating immunostimulatory effects of STING activation in tumor models [45], viral infections [46] and as a vaccine adjuvant application [47]. To our knowledge, however, this is the first study showing that activation of the STING pathway can restore innate immune activation in the context of a staphylococcal biofilm environment. In the presence of BCM, addition of 2′3′-cGAMP not only restored downstream interferon signaling but also markedly enhanced innate immune activation markers, including TLR2, MHC class II, and the co-stimulatory molecules CD80 and CD86. Interestingly, when cells were first exposed to BCM followed by delayed 2′3′-cGAMP stimulation, the enhanced immune activation observed under simultaneous treatment was no longer evident. This finding suggests a synergistic interaction between BCM-derived signals and STING activation that requires concurrent stimulation and is lost upon sequential exposure. Combinatory stimulation of immune cells with STING agonists and CpG ODN was reported to up-regulate costimulatory markers MHC II and CD86 as well as proinflammatory cytokines suggesting a synergistic effect via a TRAF6-dependent process [48]. A comparable phenomenon has been described in a tumor study, where cGAMP enhanced CpG-induced TLR9-mediated immune activation, leading to an improved anti-tumor response [49]. In the context of staphylococcal biofilms, where pattern recognition receptor signaling such as TLR2 activation is dampened [18, 50, 51], pharmacological activation of STING may therefore represent a strategy to re-establish or boost antibacterial immunity.

### Strengths and limitations

A limitation of this study is the predominant use of the murine macrophage cell line RAW 264.7, in which intrinsic alterations in innate immune signaling cannot be fully excluded and may influence the responsiveness of the cGAS-STING pathway. However, key experiments performed in primary BMDMs yielded comparable results, supporting the general validity of our observations beyond this transformed cell line. Another limitation is that the precise molecular mechanism underlying the impaired activation of the cGAS-STING pathway in the SA biofilm environment could not be fully elucidated in the present study. However, our findings extend previous observations and provide key mechanistic insights, demonstrating that impaired signaling is not primarily driven by extracellular nuclease activity or degradation of the cGAS product, but instead occurs upstream of STING at the level of ligand availability and cGAS activity. While direct assessment of cGAS activity was beyond the scope of this study, our data suggest that alterations in cGAS availability or function may contribute to the observed impairment. Nevertheless, the biofilm-derived factor(s) responsible for this effect and the exact regulatory mechanisms remain to be defined. Small metabolites accumulating in the biofilm environment, particularly lactate, may represent plausible candidates capable of modulating cGAS activity through metabolite-driven post-translational modifications such as protein lactylation. Importantly, despite this unresolved mechanism, our findings consistently show that direct activation of STING by exogenous 2′3′-cGAMP is sufficient to restore and potentiate immune signaling. This observation highlights the functional relevance of the identified impairment and further supports the concept that targeting STING downstream of cGAS may represent a potential strategy to overcome biofilm-associated immune suppression. Taken together, these findings provide new insights into the interaction between SA biofilms and host DNA sensing pathways and point towards therapeutic approaches aimed at restoring innate immune activation and an effective host defense in chronic implant-related bone infections.

## 6. Author contribution

ES conceived and designed the study, acquired, analyzed and interpreted the data and drafted the manuscript. CEPC performed experiments and contributed to data acquisition and analysis. RS participated in experiments and data acquisition. KFK contributed to data interpretation and critically revised the manuscript. All authors read and approved the final manuscript.

## 7. Acknowledgements

We would like to thank Magdalena Urlaub and Ann-Katrin Niedermayer for help with the experiments and Gabriele Sonnenmoser for technical assistance. Furthermore, we thank Dr. Lan-Sun Chen and Dr. Vivek V. Thacker for valuable discussion. This study did not receive external funding.

## 8. Declaration of generative AI and AI-assisted technologies in the manuscript preparation process

During the preparation of this work ChatGPT was used for English language editing and Elicit was used to assist with literature discovery. After using this tool/service, the authors reviewed and edited the content as needed and take full responsibility for the content of the published article.

## 9. Data availability statement

The data that support the findings of this study are available from the corresponding author upon reasonable request.

## Supplementary Material

**Suppl. Figure 1.**
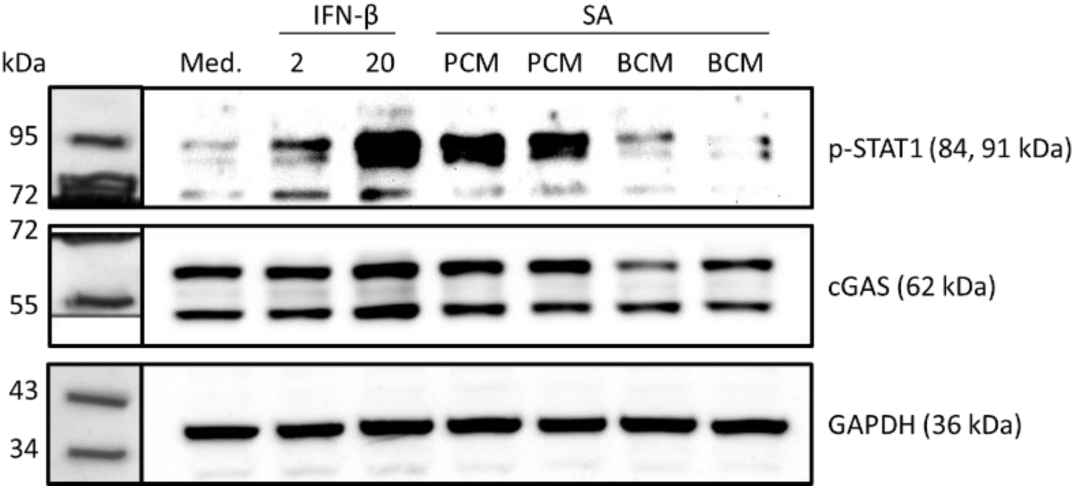
Full blot of cropped WB image of Fig. 2B is shown. RAW 264.7 cells were treated with recombinant mouse IFN-β (2 or 20 ng/ml) and SA PCM or BCM of two batches for 20 hours. Protein content (cGAS) and IFN pathway activation (p-STAT1) was assessed with WB. GAPDH was used as loading control of 10 µg protein per lane.

**Suppl. Figure 2.**
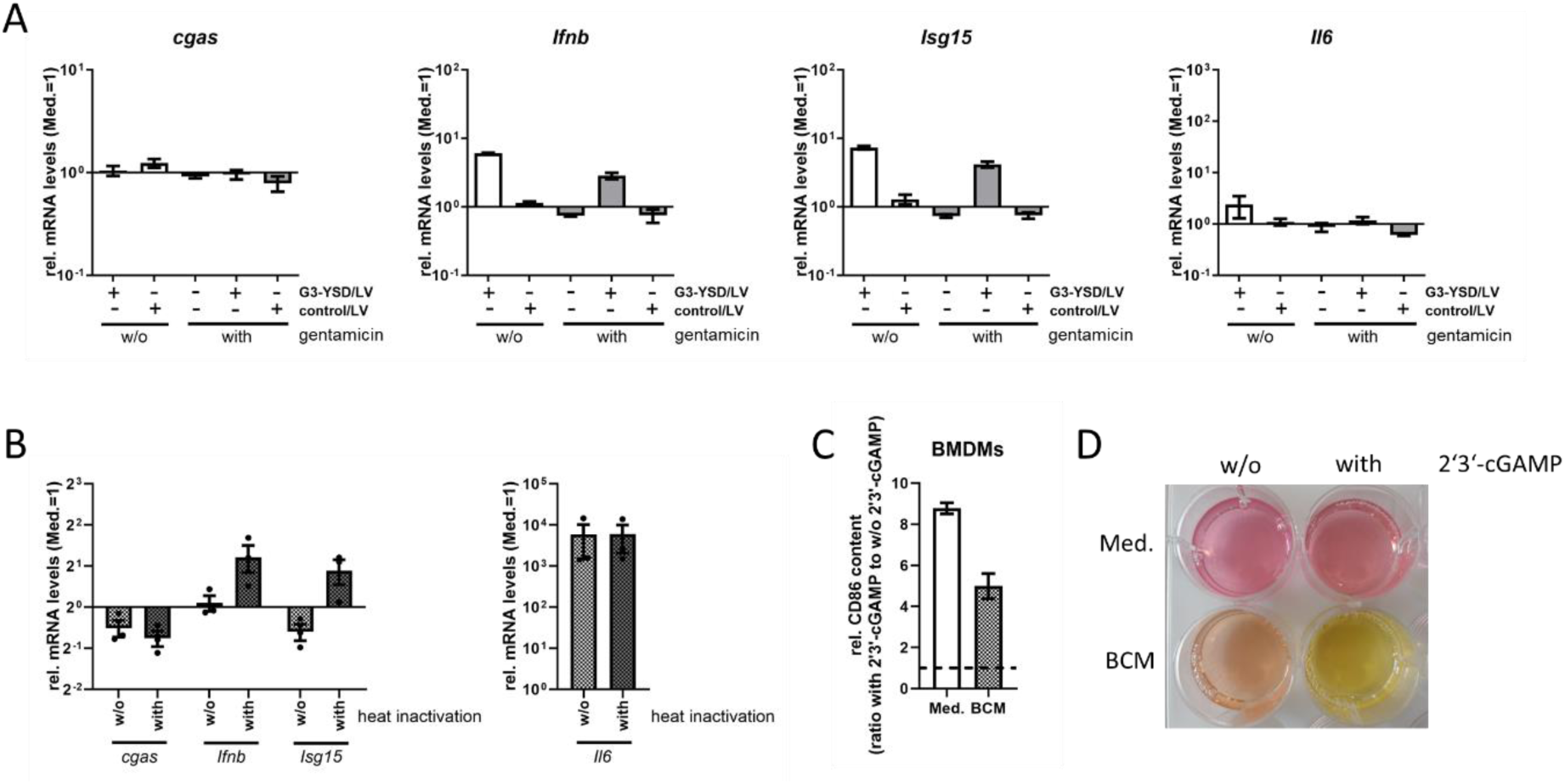
A) Effect of gentamicin on cGAS activation. RAW 264.7 cells were stimulated with G3-YSD/LyoVec^TM^ for 20 h with or without the addition of 100 µg/ml gentamicin in the medium. Induction of a type I IFN response and general immune activation was assessed by RT-qPCR analysis of *cgas*, *Ifnb*, *Isg15* and *Il6*. mRNA levels were normalized to the reference gene *Hprt1* and expressed relative to the medium-treated control (Med.). Graphs show mean ± SEM from two independent experiments. B) Effect of heat-inactivation of SA BCM on macrophage immune activation. Induction of a type I IFN response and general immune activation was assessed by RT-qPCR analysis of *cgas*, *Ifnb*, *Isg15* and *Il6*. mRNA levels were normalized to the reference gene *Hprt1* and expressed relative to the medium-treated control (Med.). Graphs show mean ± SEM from three independent experiments. C+D) Effect of 2’3’-cGAMP on immune activation of primary macrophages (BMDMs) by SA BCM. BMDMs were treated simultaneously with SA BCM and 2’3’-cGAMP for 24 hours and immune activation was assessed. C shows protein content of surface marker CD86 as ratio with to without 2’3’-cGAMP in medium (Med.) or BCM. Graphs show mean ± SEM from n=2 mice. D shows appearance of media after stimulation of BMDMs with SA BCM ± 2’3’-cGAMP.

